# NCLX controls hepatic mitochondrial Ca^2+^ extrusion and couples hormone-mediated mitochondrial Ca^2+^ oscillations with gluconeogenesis

**DOI:** 10.1101/2024.02.09.579606

**Authors:** Mahmoud Taha, Essam A. Assali, Grace E. Stuzmann, Orian S. Shirihai, Michal Hershfinkel, Israel Sekler

## Abstract

Hepatic Ca^2+^ signaling is emerging as a key factor in mediating gluconeogenesis. However, the identity of the hepatic mitochondrial Ca^2+^ transporter is controversial and the role of mitochondria in controlling hormonal Ca^2+^ signaling and linking them to metabolic activity is poorly understood. We first interrogated the role of the mitochondrial Na^+^/Ca^2+^ exchanger NCLX by triggering cytosolic Ca^2+^ purinergic signaling in primary hepatocytes, and Ca^2+^ responses in isolated mitochondria from WT, global NCLX KO, and conditional hepatic NCLX KO mice models. We monitored a higher rate of Na^+^-dependent mitochondrial Ca^2+^ efflux in NCLX-expressing hepatocytes, indicating that it constitutes the major Ca^2+^ efflux pathway. We then asked if NCLX is controlling the hormone-dependent mitochondrial Ca^2+^ oscillations by employing physiological concentrations of glucagon and vasopressin. Consistent with previous studies, hormone applications triggered mitochondrial Ca^2+^ oscillations in WT hepatocytes. In NCLX KO hepatocytes the cytosolic oscillations persisted, however, the mitochondrial Ca^2+^ oscillations were suppressed. To further understand the metabolic role of NCLX in the hepatic system, we examined gluconeogenic function in vivo and ex vitro by monitoring hepatic glucose production. We found that blood glucose dropped faster in the conditional KO mice and their hepatic glucagon-dependent glucose production was reduced, indicating that gluconeogenesis was impaired in hepatic conditional NCLX KO mice. Taken together, our results indicate that NCLX is the primary Ca^2+^ extruder in hepatocytes and is required for mediating the hormone-dependent mitochondrial Ca^2+^ oscillations and gluconeogenesis.

**Significance:** Hepatic Ca^2+^ signaling is crucial for gluconeogenesis, but the mitochondrial control of this process is not resolved. This study identifies the mitochondrial transporter, NCLX, as a critical link between hormonal-dependent mitochondrial Ca^2+^ oscillations and gluconeogenesis. We first show that NCLX is the major hepatic mitochondrial efflux pathway. We then demonstrate that NCLX is required for glucagon-dependent mitochondrial Ca^2+^ oscillations and the acceleration of mitochondrial oxidative function. Using a conditional hepatic NCLX-null mouse model, we show that NCLX is required for maintaining hepatic glucose production during fasting and in response to glucagon stimulation. Overall, the study identifies NCLX as the integrator of hepatic mitochondrial Ca^2+^ signaling, required for gluconeogenesis.

## Introduction

Mitochondrial Ca^2+^ dynamics has a pivotal role in orchestrating cellular bioenergetics, Ca^2+^ signaling and regulation of cell death (1). Ca^2+^ enters the mitochondria via the mitochondrial Ca^2+^ uniporter (MCU) complex and is subsequently extruded either through the mitochondrial Na^+^/ Ca^2+^ exchanger, molecularly termed NCLX, or potentially through a less characterized H^+^/Ca^2+^ exchange mechanism remained to be further clarified physiologically and molecularly (2–4).

Calcium signaling has an intriguing metabolic role in the liver (5). Hepatocytes are distinguished by their abundance of contact sites between the endoplasmic reticulum (ER) and mitochondria, which facilitate localized and efficient transmission of Ca^2+^ signals from the inositol triphosphate receptors (IP_3_R) to the mitochondrial matrix (6). Seminal studies showed that hepatic hormones such as glucagon, when applied at physiological concentrations, evoke IP_3_R-dependent cytoplasmic Ca^2+^ ([Ca^2+^]c) oscillations (5–8). Hepatocytes demonstrate that each [Ca^2+^]c spike independently propagates from the cytosol to the mitochondria, where it triggers NADH/NAD+ transients (7, 9, 10) Therefore, hepatic [Ca^2+^] oscillatory waves are intrinsically coupled to metabolism flux and mitochondrial function.

The robust correlation between [Ca^2+^]m oscillations and metabolic alterations in hepatocytes, provides a tool for investigating downstream physiological pathways mediated by glucagon activity, especially during the fasting state. Incremental glucagon levels during fasting increase the glucagon-to-insulin ratio in the portal circulation, thereby enhancing the liver glucose output via glycogenolysis then gluconeogenesis. Regulated allosterically, glycogenolysis facilitates rapid mobilization and depletion of glycogen stores. Gluconeogenesis is a slower more complex process requiring transcriptional regulation and activation of intertwined pathways breaking down essential blocks such as amino acids and fatty acids, thus facilitating sustained and continuous hepatic glucose production. Initially, the role of glucagon in gluconeogenesis was predominantly ascribed to transcriptional mechanisms through the activation of cAMP-responsive element binding protein (CREB), located on the promoters of key gluconeogenic genes such as PEP-CK, pyruvate carboxylase and others (11). However, recent studies challenged the dominance of the glucagon-dependent transcriptional pathway in gluconeogenesis induction and underscored an alternative pathway involving Ca^2+^ signaling through an IP_3_R isoform, where phosphorylation of INSP_3_R1 by glucagon-activated PKA was shown to play a prominent role in hepatic glucose production (12, 13). However, it remains unknown whether the canonical mitochondrial Ca^2+^ oscillations contribute to glucagon-dependent gluconeogenesis. Furthermore, the distinct roles of the mitochondrial Ca^2+^ transporters in hepatic Ca^2+^ signaling and in modulating gluconeogenesis remains to be addressed.

A major challenge in deciphering the role of mitochondrial Ca^2+^ signaling in gluconeogenesis is the unresolved molecular identity for the hepatic Ca^2+^ efflux pathway. Earlier studies, mostly carried out on isolated liver mitochondria, proposed that hepatic mitochondrial Ca^2+^ removal is mediated by a Na^+^-independent mechanism, presumably H^+^/Ca^2+^-dependent exchange pathway (14). However, other studies utilizing isolated mitochondria from mice pre-treated with norepinephrine (NE) or other PKA activators, demonstrated a significant component of Na^+^-dependent mitochondrial Ca^2+^ efflux (15–19).

In this study, by employing models of global and conditional hepatic-specific knockout (KO) for NCLX, we show that NCLX has a critical role facilitating mitochondrial Ca^2+^ efflux in liver hepatocytes. Furthermore, our data reveal a dominant Na^+^-dependent Ca^2+^ extrusion mode in isolated mitochondria as well as permeabilized and intact hepatocytes. Surprisingly, the hepatic ablation of NCLX completely abolished the induction of mitochondrial Ca^2+^ oscillations by hepatic hormones, such as glucagon and vasopressin. Lastly, our findings elucidate a crucial role of NCLX in mediating glucagon-dependent hepatic glucose production during fasting.

Remarkably, loss of NCLX inhibited gluconeogenesis tested in hepatocytes and mice, further indicating an indispensable Ca^2+^ signaling and metabolic role for NCLX activity in vitro and in vivo.

## Results

### NCLX is essential for mitochondrial Ca^2+^ efflux in hepatocytes

To study the role of NCLX in mitochondrial Ca^2+^ efflux in hepatocytes, we first used a conditional hepatocyte-specific NCLX KO (KO) utilizing the Cre-loxP recombination system for gene targeting (20, 21). Mice with loxP-flanked NCLX alleles (denoted as NCLX^fl/fl^) were tail-vein injected with liver-tropic AAV8 viral vector containing a hepatocyte-specific thyroid-binding globulin (TBG) promoter driving a Cre recombinase expression, or its inactive mutant Cre recombinase that served as control (**Fig. 1A**). To determine if the Cre virus effectively knocked down NCLX, we conducted a western blot analysis, which confirmed ablation of NCLX in the liver tissue, while maintaining the normal differential distribution of NCLX protein levels in other tissues (**Fig. 1B**). Additionally, no alteration in MCU expression was detected in the surveyed tissues (**Fig. 1C**). To study the contribution of NCLX to hepatic mitochondrial Ca^2+^ extrusion, primary hepatocytes from conditional KO (cKO) mice and their respective controls were isolated and stained with the mitochondrial Ca^2+^ reporter, Rhod-2 AM. We then induced mitochondrial Ca^2+^ transients by applying ATP-dependent purinergic intracellular Ca^2+^ rise and subsequent mitochondrial Ca^2+^ fluxes (**Fig. 1D**). Our results show that in hepatocytes lacking NCLX mitochondrial Ca^2+^ influx and uptake were unaffected. In contrast, we find a ∼ 2.5-fold slower mitochondrial Ca^2+^ efflux in NCLX cKO compared to control hepatocytes (**Fig. 1E,F**). Additionally, monitoring cytosolic Ca^2+^ transients using a Flou4-AM, revealed no alterations in cytosolic Ca^2+^ kinetics or amplitude in primary hepatocytes lacking NCLX (**Fig. 1G,H**).

**Fig. 1.**
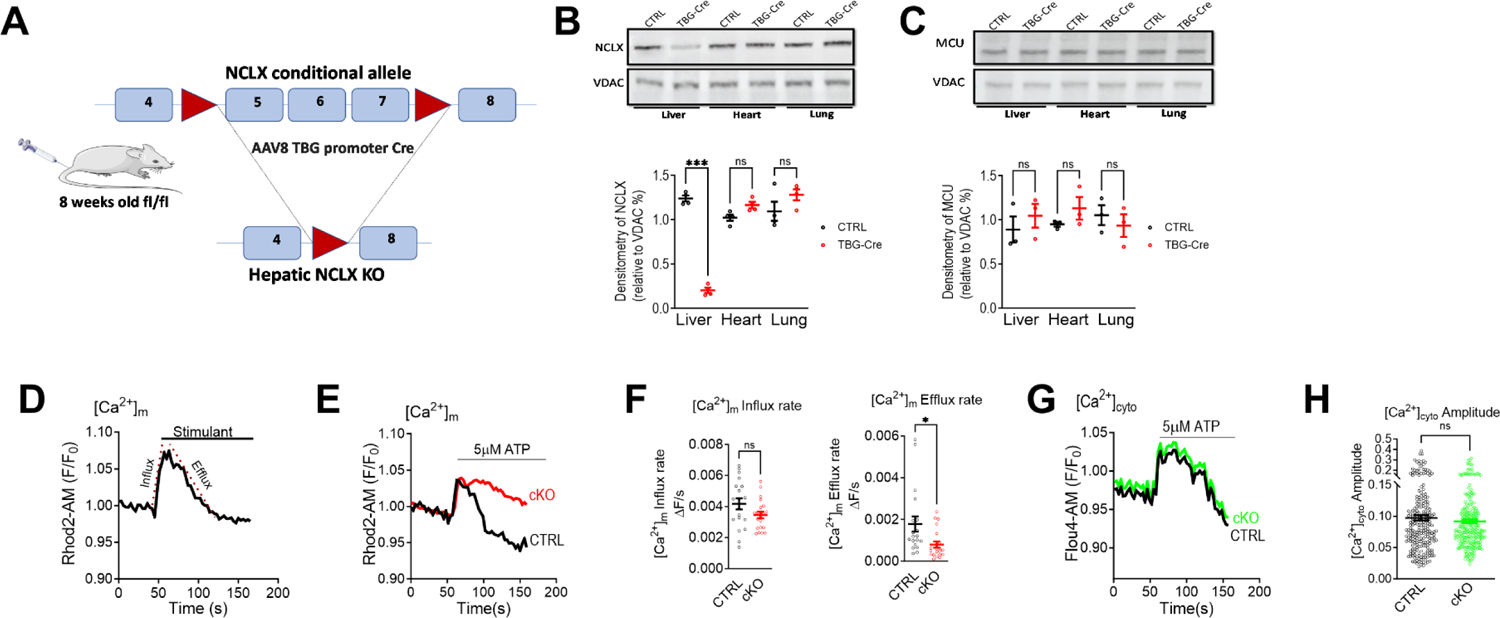
NCLX is essential for mitochondrial Ca^2+^ efflux in hepatocytes. (A) Schematic strategy of conditional hepatic NCLX knockout: *(top panel)* Map of *NCLX* gene (slc8b1) with exons 5-7 flanked by two loxP sites. *(bottom panel)* Mice were injected via the tail vein with an AAV8-Cre under the control of a hepatocyte-specific promoter, thyroid-binding globulin (TBG) with subsequent recombination leading to the truncation of the NCLX gene in the hepatocytes only. AAV8-TBG-Null-injected littermates were used as controls. (B) Western blot *(top panel)* and densitometry analysis *(bottom panel)* of NCLX expression in tissues obtained from mice injected either with a hepatic viral Cre vector (TBG-CRE) or TBG-Null (CTRL). VDAC1 was used as a loading control (N = 4 mice per group). (C) Western blot *(top panel)* and densitometry analysis *(bottom panel)* of MCU expression in tissues obtained from mice injected with either hepatic viral Cre vector (TBG-CRE) or TBG-Null (CTRL). VDAC1 was used as loading control (N = 3 mice per group). (D-F) Mitochondrial Ca^2+^ kinetics evoked by ATP in cultured primary hepatocytes isolated from NCLX conditional KO (cKO) mice (cKO, Red) and their controls (CTRL, black). (D) Representative fluorescence transients in primary hepatocytes loaded with the mitochondrial Ca^2+^ dye, Rhod2-AM, application of a stimulus (ATP) at the indicated time and [Ca^2+^]m was monitored. The dashed lines represent the linear fit used to calculate the [Ca^2+^]m influx and efflux rates. (E) Representative [Ca^2+^]m transients of NCLX cKO hepatocytes and their controls (CTRL), triggered by ATP (5μM). (F) Quantification of [Ca^2+^]m influx and efflux rates for CTRL (n = 19) and cKO (n = 21). (G,H) Cytosolic Ca^2+^ kinetics evoked by ATP in cultured primary hepatocytes isolated from NCLX cKO (cKO, Green) and their controls (CTRL, black). (G) Representative cytosolic Ca^2+^ [Ca^2+^]c transients of cKO hepatocytes, loaded with Flou4-AM, and their controls, triggered by ATP (5μM). (H) Quantification of [Ca^2+^]c amplitude for CTRL (n = 235) and cKO (n = 244). All values are represented as mean ± SEM, \**p* < 0.05; \*\**p* < 0.01; \*\*\**p* < 0.0001, N.S-non-significant (two-tailed unpaired *t*-test for comparisons between two groups was used and one-way ANOVA with Tukey’s multiple comparison test was used for three or more groups).

We extended our examination to an additional mouse model with global deletion of NCLX, using the same experimental settings to monitor mitochondrial and cytosolic Ca^2+^ upon purinergic stimulation. Similarly to results shown in figure 1, the mitochondrial Ca^2+^ efflux was compromised in hepatocytes with NCLX deletion, yet Ca^2+^ influx rates were unaffected **(Fig. S1A-C)** with no alterations in cytosolic Ca^2+^ kinetics **(Fig. S1E,F)**. Altogether, the results of this set of experiments indicate that NCLX plays a key role in facilitating hepatic mitochondrial Ca^2+^ extrusion.

### NCLX mediates a mitochondrial Na^+^-dependent Ca^2+^ efflux in hepatocytes

The understanding of the ion-dependence mode essential for mitochondrial Ca^2+^ release in hepatocytes has faced significant challenges, primarily arising from conflicting reports of Na^+^-dependent versus Na^+^-independent mechanisms, predominantly from studies done on isolated liver mitochondria(14, 22). To study the monovalent-cation dependence required for hepatic mitochondrial Ca^2+^ extrusion, we employed several approaches. First, we utilized a setup where we loaded primary hepatocytes with Rhod2-AM, then permeabilized the cell membrane by digitonin to facilitate ion permeability, and immediately monitored mitochondrial Ca^2+^. Ionic selectivity required for mitochondrial Ca^2+^ efflux was assessed using either Na^+^, or NMDG^+^ as control, or Li^+^ that is uniquely transported by NCLX (4) (**Fig. 2A**). We initially introduced a Ca^2+^ bolus which resulted in rapid Ca^2+^ uptake, then we switched to Ca^2+^-free solution containing one of the three ions, all applied at the same concentration. Ruthenium Red (RR), which blocks the uniporter, was present in all Ca^2+^-free solutions (**Fig. 2B**). Notably, the presence of extra-mitochondrial Na^+^ and Li^+^ activated mitochondrial Ca^2+^ efflux in WT hepatocytes (by ∼ 1.4-fold with Li^+^ and by ∼ 1.6-fold with Na^+^ compared to NMDG^+^). Conversely, NCLX KO hepatocytes did not demonstrate any significant Na^+^ or Li^+^-dependent mitochondrial Ca^2+^ efflux when compared to NMDG^+^ (**Fig. 2C**). Second, we explored the mode of Ca^2+^ extrusion and the molecular role of NCLX in liver isolated mitochondria (22) (**Fig. 2D**). We applied an indirect mitochondrial fluorescent Ca^2+^ monitoring assay that overcomes mitochondrial Ca^2+^ buffering issues using the impermeable acid form of the low affinity fluorescent Ca^2+^ sensor Oregon Green. We maintained harvested liver mitochondria in an intracellular-mimicking solution with or without Na^+^ (NMDG^+^ isosmotically replaced Na^+^) (23–25). Initially, a Ca^2+^ bolus was added to evoke mitochondrial Ca^2+^ uptake, followed by an addition of Ca^2+^-free solution without or with Na^+^ to activate the exchanger and subsequent increase in the rate of mitochondrial Ca^2+^ removal reflected by the fluorescence rise was monitored (**Fig. 2E**). As in the previous experimental setup, mitochondrial Ca^2+^ efflux demonstrated a strong dependence on the presence of Na^+^ in the intracellular-mimicking solution and was enhanced by ∼ 3-fold compared to Na^+^-free conditions. In contrast, NCLX KO mitochondria displayed a significant reduction in Ca^2+^ extrusion, with no observed Na^+^-dependence (**Fig. 2F**). Finally, to further interrogate whether Ca^2+^ efflux is Na^+^ dependent in human hepatocytes, we utilized HepG2 cells. During Rhod2-AM imaging, Hep2G cells were treated with ATP while maintained in media containing Na^+^, Li^+^ or NMDG^+^. Our results show that Na^+^ and Li^+^ activated the mitochondrial Ca^2+^ efflux by ∼ 3.5-fold and ∼ 2-fold increase, respectively, thereby these results further support an NCLX activity in hepatocytes **(Fig. S2A-C)**. Altogether, the results of this part indicate that in hepatocytes the major Ca^2+^ efflux pathway is predominantly Na^+^-dependent and mediated by NCLX.

**Fig. 2.**
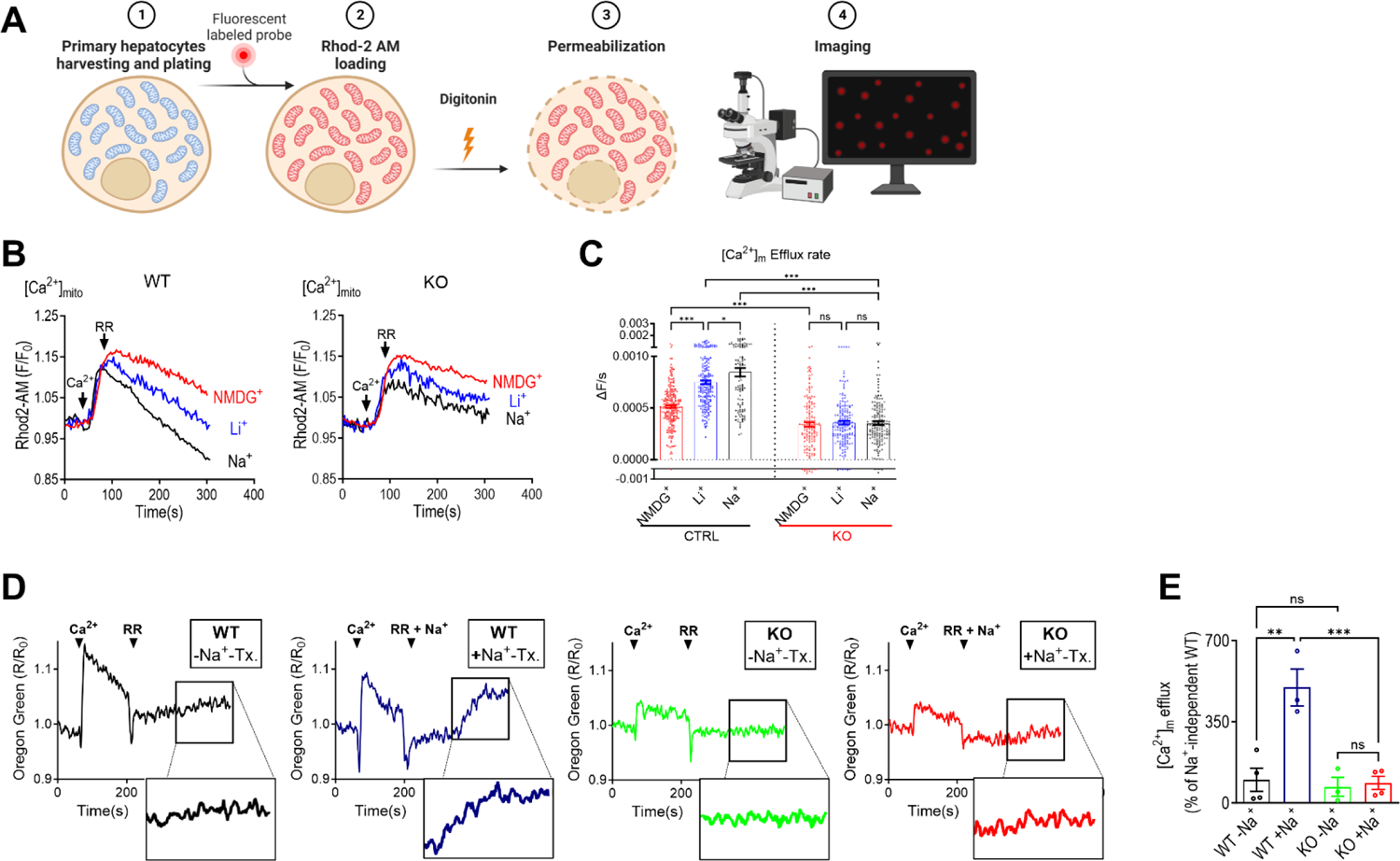
NCLX mediates Na^+^-dependent mitochondrial Ca^2+^ efflux in hepatocytes. (A-C) Analysis of monovalent cation-dependence of [Ca^2+^]m efflux in permeabilized hepatocytes. (A) Schematic cartoon depicting [Ca^2+^]m monitoring in permeabilized primary hepatocytes. (B) Representative traces of [Ca^2+^]m transients in primary WT *(left panel)* and NCLX KO permeabilized hepatocytes *(right panel)*. Ca^2+^ (6 µM) was applied to permeabilized cells and subsequently 10 µM ruthenium red (RR) was applied (at the indicated times). Mitochondrial Ca^2+^ extrusion was monitored in the presence of either 10 mM Na^+^, Li^+^ or NMDG^+^ (used as a cationic replacement of Na^+^). Detailed experimental design is described in Methods section. (C) Quantification [Ca^2+^]m efflux rates. WT hepatocytes: (n = 226 for NMDG^+^, n = 197 for Li^+^ and n = 109 for Na^+^); in NCLX-KO hepatocytes: (n = 153 for NMDG^+^, n = 172 for Li^+^ and n = 171 for Na^+^). (D-E) Mitochondrial calcium analysis of isolated mouse liver mitochondria. (D) Representative fluorescent traces of extra-mitochondrial Ca^2+^ transients monitored in isolated mitochondria from liver of WT or NCLX KO mice in the presence of Oregon-Green (300nM), triggered by the addition of 6μM free Ca^2+^ followed by the administration of either Na^+^ or NMDG^+^ (Na^+^-free) solutions supplemented with 10µM RR. Decrease of extra-mitochondrial Ca^2+^ (fluorescence signal) is representative of [Ca^2+^]m uptake while an increase in fluorescence signal is indicative of [Ca^2+^]m efflux. Tx, treatment. (E) Quantification of [Ca^2+^]m efflux rates shown in WT: (n = 4 for Na^+^-free, n= 3 for Na^+^-containing), in NCLX-KO hepatocytes: (n = 3 for Na^+^-free, n= 4 for Na^+^-containing). All values are represented as mean ± SEM, \**p* < 0.05; \*\**p* < 0.01; \*\*\**p* < 0.0001, N.S-non-significant (two-tailed unpaired *t*-test for comparisons between two groups was used and one-way ANOVA with Tukey’s multiple comparison test was used for three or more groups).

### NCLX is indispensable for vasopressin and glucagon-induced oscillations of mitochondrial Ca^2+^

Hepatic hormones, most prominently glucagon and vasopressin, at physiological concentrations elicit intracellular Ca^2+^ oscillations which are then propagated individually into the mitochondrial matrix (10, 26). These mitochondrial Ca^2+^ oscillations have a key role in tuning the mitochondrial redox state (7, 8, 10). Mechanistically, these oscillatory waves are largely dependent on the IP_3_R signaling system, yet the mitochondrial players orchestrating these oscillations are not well-characterized (7, 9). Consequently, given the crucial function of NCLX in controlling mitochondrial efflux within the hepatocytes, we sought to study its role in response to glucagon stimuli, which was applied at physiological concentrations to hepatocytes isolated from NCLX cKO mice and their respective controls. Remarkably, while control hepatocytes responded to glucagon with the expected low-frequency intra-cellular and mitochondrial Ca^2+^ oscillations (**Fig. 3A,C**), NCLX cKO hepatocytes exhibited a complete cessation of glucagon-dependent mitochondrial Ca^2+^ oscillations (**Fig. 3A,B**). Intracellular Ca^2+^ oscillations persisted in the NCLX cKO hepatocytes, without an impact on their frequency and showed a modest decrease in area under the curve of the cytosolic Ca^2+^ responses between the NCLX cKO and WT hepatocytes (**Fig. 3C,D and Fig. S3A,B)**. Next, we broadened our analysis to include vasopressin (VP) signaling (9, 26), which at physiological concentrations as low as 1nM triggers intracellular Ca^2+^ oscillations akin to glucagon. Application of VP to WT hepatocytes incited mitochondrial and intracellular Ca^2+^ oscillations, as anticipated (**Fig. 3E,G**). In contrast, VP-failed to evoke mitochondrial Ca^2+^ oscillations in NCLX cKO hepatocytes, similar to glucagon and thus further validating the essential role of NCLX in mediating mitochondrial Ca^2+^ oscillations (**Fig. 3E,F**). Again, intracellular Ca^2+^ oscillations and their frequency in the cKO remained largely unaffected when compared to the WT, with a modest decrease in the area under the curve of the individual spikes (**Fig. 3G,H and Fig. S3C,D)**.

**Fig. 3.**
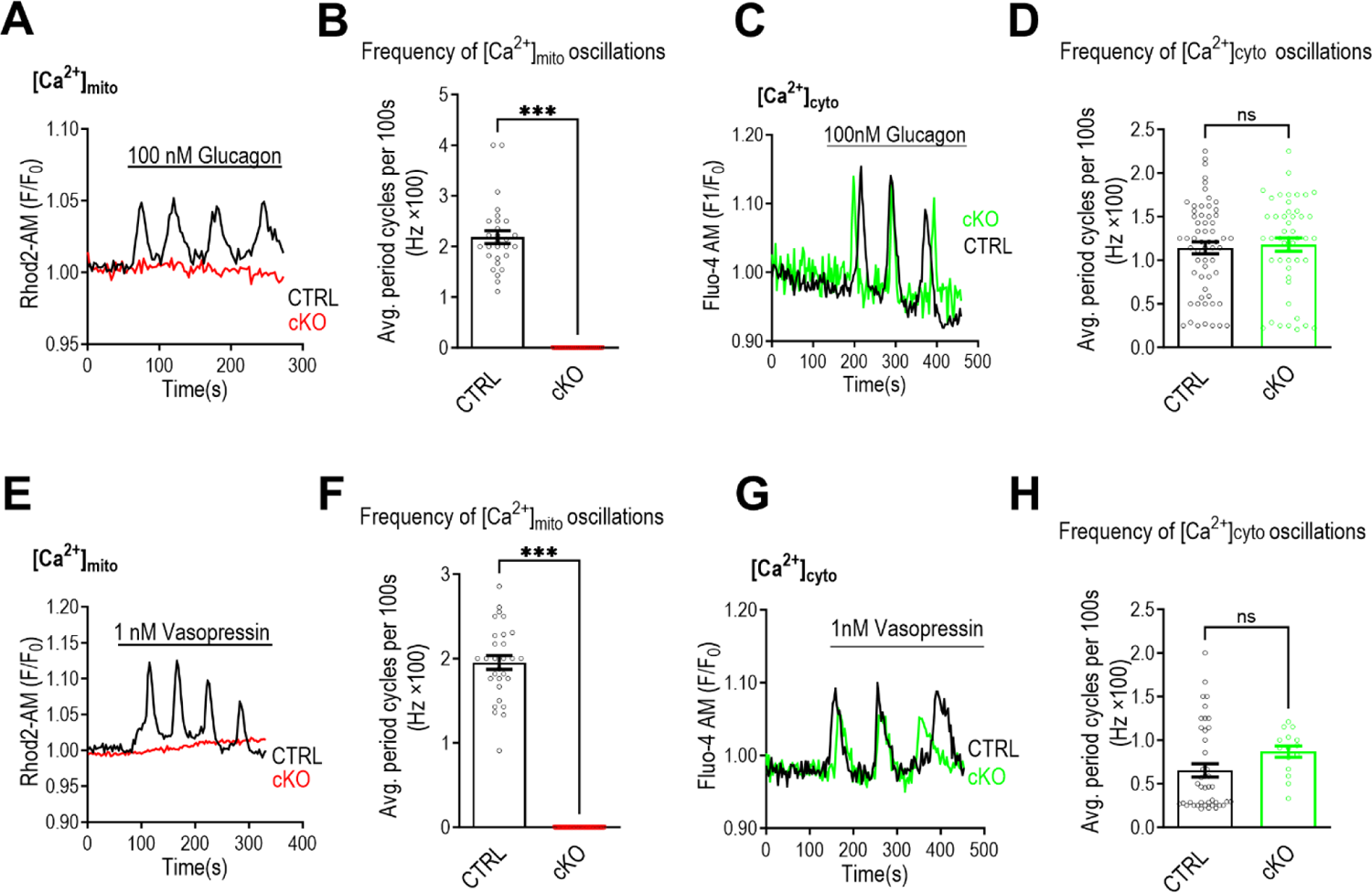
NCLX is required for Ca^2+^ oscillations generated by physiological concentrations of glucagon and vasopressin. (A,B) Mitochondrial Ca^2+^ oscillations triggered by physiological concentrations of glucagon in cultured primary hepatocytes isolated from NCLX cKO mice (cKO, Red) and their controls (CTRL, black). [Ca^2+^]m was monitored by Rhod2-AM. (A) Representative fluorescence traces of [Ca^2+^]m, triggered by glucagon (100nM) in NCLX cKO hepatocytes and their controls (CTRL). (B) Quantification of [Ca^2+^]m frequency of oscillations shown in (A) for CTRL (n = 28) and NCLX cKO hepatocytes (n = 29). (C,D) Cytosolic Ca^2+^ oscillations triggered by physiological concentrations of glucagon in cultured primary hepatocytes isolated from NCLX cKO mice (cKO, Green) and their controls (CTRL, black). Cytosolic Ca^2+^ was monitored by Flou4-AM. (C) Representative fluorescence traces of cytosolic Ca^2+^ oscillations triggered by glucagon (100 nM) in NCLX cKO hepatocytes and their controls (CTRL). (D) Quantification of [Ca^2+^]c frequency of oscillations shown in (C) for CTRL (n = 60) and NCLX cKO hepatocytes (n = 48). (E,F) Mitochondrial Ca^2+^ oscillations triggered by physiological concentrations of vasopressin (VP) in cultured primary hepatocytes isolated from NCLX cKO mice (cKO, Red) and their controls (CTRL, black). Mitochondrial Ca^2+^ was monitored by Rhod2-AM. (E) Representative fluorescence traces of mitochondrial Ca^2+^ oscillations triggered by VP (1nM) in NCLX cKO hepatocytes and their controls (CTRL). (F) Quantification of mitochondrial Ca^2+^ frequency of oscillations shown in (G) for CTRL (n = 29) and NCLX cKO hepatocytes (n = 42). (G,H) Cytosolic Ca^2+^ oscillations triggered by physiological concentrations of VP in cultured primary hepatocytes isolated from NCLX cKO mice (cKO, Green) and their controls (CTRL, black). Cytosolic Ca^2+^ was monitored by Flou4-AM. (G) Representative fluorescence traces of cytosolic Ca^2+^ oscillations triggered by VP (1 nM) in NCLX cKO hepatocytes and their controls (CTRL). (H) Quantification of [Ca^2+^]c frequency of oscillations shown in (A) for CTRL (n = 42) and NCLX cKO hepatocytes (n = 15). All values are represented as mean ± SEM, \**p* < 0.05; \*\**p* < 0.01; \*\*\*\**p* < 0.0001, N.S-non-significant (two-tailed unpaired *t*-test for comparisons between two groups was used).

This set of experiments was then performed in a hepatocytes derived from global NCLX KO mice and their WT controls, consistently demonstrated that while glucagon and VP triggered intracellular and mitochondrial Ca^2+^ oscillations in WT hepatocytes, neither hormone could induce mitochondrial Ca^2+^ oscillations in the NCLX KO hepatocytes **(Fig. S4A-B, G-H)**.

Intracellular Ca^2+^ oscillations were largely preserved across both genotypes and no disparity in the magnitude of cytosolic Ca^2+^ responses was noted, albeit with a modest decrease of cytosolic Ca^2+^ oscillation frequency only with VP in the global NCLX KO hepatocytes **(Fig. S4A-L)**.

In summary, these findings show that NCLX is a key regulator for mitochondrial Ca^2+^ oscillations evoked by hepatic hormones underscoring the physiological role NCLX in hepatic mitochondrial calcium signaling.

### NCLX is essential for glucagon-mediated hepatic gluconeogenesis

In light of the significant role of NCLX in facilitating glucagon-dependent mitochondrial Ca^2+^ oscillations, we hypothesized that NCLX activity is essential for glucagon-dependent gluconeogenesis. To investigate the potential role of NCLX in hepatic glucose production (HGP), we first continuously monitored blood glucose levels in hepatic NCLX cKO mice and their control littermates under fasting conditions **(FIG 4A)**. Interestingly, while no discernible difference was observed during the initial 8 hours (which corresponds to the time required to deplete glycogen stores), a subsequent steady fast decline in blood glucose levels was noted in the NCLX cKO mice. This decline continued until their blood glucose levels reached hypoglycemic levels, defined by a cut-off of 60 mg/dl, in contrast to the control mice, which maintained significantly higher blood glucose concentrations **(FIG. 4A)**.

Next, to confirm that HGP primarily arises from gluconeogenesis, we assessed the activity rate of hepatic pyruvate carboxylase (PC), an essential enzyme and a rate controlling factor for gluconeogenesis process(11, 13, 27). Utilizing liver homogenates isolated from fasted and fed NCLX cKO and their controls, an increase in PC activity rate was noted in the fasted state of control mice while the NCLX cKO mice failed to increase PC activity upon fasting (**Fig. 4B**).

**Fig. 4.**
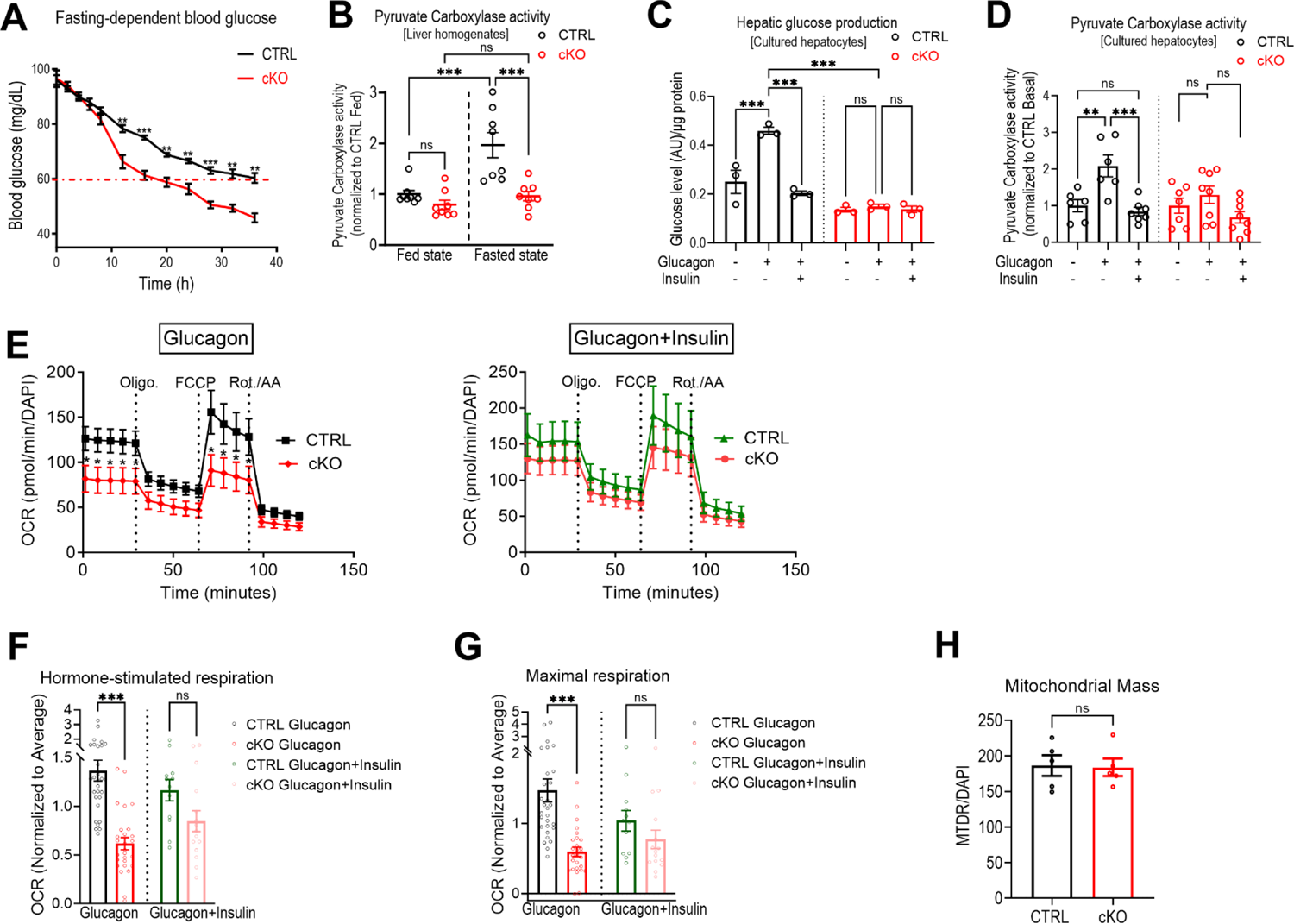
NCLX is required for glucagon-mediated hepatic glucose production. (A) Blood glucose measurements at indicated time intervals in mice during fasting from NCLX cKO (cKO) and CTRL mice (n=4 each). The dashed red line indicates the lower threshold of physiological glycemic control (60 mg/dL). (B) Hepatic pyruvate carboxylase activity determined in liver homogenates from - NCLX cKO and CTRL in fed and fasted conditions (n = 8 for each condition). (C) Gluconeogenesis assessed by glucose production in response to lactate and pyruvate in cultured primary mouse hepatocytes isolated from NCLX cKO and CTRL mice. Glucagon (100 nM) was used to induce gluconeogenesis. Co-adminstration of insulin (100 nM) with glucagon was used as a negative control (n = 3 for each condition). (D) Hepatic pyruvate carboxylase activity assessed in response to lactate and pyruvate in cultured primary mouse hepatocytes from NCLX cKO and CTRL mice (For basal enzymatic activity n = 6 CTRL and n = 7 cKO; For glucagon stimulation n = 6 CTRL and n = 8 cKO; for co-stimulation of insulin and glucagon n = 7 CTRL and n = 8 cKO). (E-H) Oxygen consumption rates (OCR) of NCLX cKO and CTRL primary hepatocytes pretreated either with glucagon alone *(left panel)* or co-treated with insulin and glucagon *(right panel)*. (E) Representative OCR traces of NCLX cKO and CTRL primary hepatocytes pretreated with glucagon *(left panel)* or co-treated with insulin and glucagon *(right panel)*. Oxygen consumption was measured in real-time under basal conditions and in response to indicated mitochondrial modulating compounds. (F) Quantification of hormonal stimulatory response in primary hepatocytes, for glucagon stimulation n = 29 CTRL and n = 27 NCLX cKO; for insulin and glucagon co-stimulation n = 12 CTRL and n = 15 NCLX cKO. (G) Quantification of maximal respiration induced by FCCP in primary hepatocytes, for glucagon stimulation n = 29 CTRL and n = 27 NCLX cKO; for insulin and glucagon co-stimulation n = 12 CTRL and n = 15 NCLX cKO (CRE). (H) Mitochondrial mass assessed by MTDR staining in plated primary hepatocytes (n = 5 for each genotype). All values are represented as mean ± SEM, \**p* < 0.05; \*\**p* < 0.01; \*\*\*\**p* < 0.0001, N.S-non-significant (two-tailed unpaired *t*-test for comparisons between two groups was used and one-way ANOVA with Tukey’s multiple comparison test was used for three or more groups).

Prolonged fasting can lead to a shift in hormonal balance, potentially affecting the gluconeogenic function of hepatocytes through altered hormonal balance(28). To determine the role of glucagon on hepatic glucose production (HGP), we evaluated glucagon-induced glucose production in cultured hepatocytes isolated from NCLX cKO mice and their control littermates **(FIG. 3C)**. Consistent with the in vivo study done in fasted mice, glucagon enhanced glucose production in cultured hepatocytes isolated from control mice. The addition of insulin to glucagon, however, blunted the rise. In contrast to the control hepatocytes, glucagon failed to stimulate glucose production in NCLX cKO hepatocytes and resulted in glucose levels similar to those produced from unstimulated hepatocytes or those co-stimulated with insulin and glucagon. We then measured PC activity in cultured primary hepatocytes. As anticipated, glucagon stimulation led to an increase in the enzyme activity rate compared to baseline or to concurrent insulin and glucagon co-stimulation, in which insulin blunted glucagon effect **(FIG. 3D)**. However, in NCLX cKO hepatocytes, no significant differences were observed in PC activity following stimulation with glucagon alone in comparison to baseline and co-stimulation with insulin and glucagon **(FIG. 3D)**.

Altogether, these in vivo and in vitro results indicate that NCLX plays a critical role in glucagon-mediated hepatic glucose production by controlling the glucagon-dependent mitochondrial Ca^2+^ signaling.

To elucidate the metabolic role of NCLX in mediating glucagon induced mitochondrial Ca^2+^ oscillations and hepatic gluconeogenic function, we conducted a seahorse respirometry analysis. This experimental approach was applied to isolated hepatocytes originating from conditional NCLX KO and their control littermates **(FIG. 4D)**. Upon subjecting the hepatocytes to glucagon pre-treatment, we observed substantial impairments in both basal and maximal respiratory capacities in the conditional NCLX KO hepatocytes **(FIG. 4E,F)**. Staining both genotypes with a mitochondrial dye (MTDR) did not reveal a discernable change in mitochondrial mass **(FIG. 4G)**. Thus, the bio-energetic findings are indicative of an impact on mitochondrial function and suggests a potential association between NCLX and the regulation of hepatic metabolism in response to glucagon stimulation. Conversely, pretreatment of insulin and glucagon did not reveal any significant differences when comparing the two genotypes (**FIG. 4E-G**) indicating that the metabolic status of NCLX KO hepatocytes remains largely intact, but their glucagon-dependent gluconeogenesis and related pathways are selectively impaired. The absence of observable alterations in insulin response between the genotypes underscores the specificity and selectivity of NCLX in hepatic metabolic hormonal homeostasis regulation.

Altogether, this set of in vivo and in vitro results indicate that NCLX plays a critical role in mediating hepatic glucose production by controlling glucagon-dependent mitochondrial Ca^2+^ signaling.

## Discussion

The molecular identity accountable for mitochondrial Ca^2+^ extrusion in the hepatocyte has been long debated and multiple mechanisms have been suggested. Our study demonstrates for the first time that NCLX is indispensable for mitochondrial calcium extrusion in hepatocytes. Furthermore, our results indicate that a Na^+^-dependent activity is the major route for mitochondrial Ca^2+^ efflux and that the presence of extracellular Na^+^ is required to fully activate Ca^2+^ extrusion. We used intact or permeabilized cells, as well as isolated mitochondria and tested human hepatocyte cell line as well as two mice models, a global NCLX KO and a conditional hepatic NCLX KO system. Notably, the presence of Na^+^ in cells and mitochondrial preparations, enhanced mitochondrial Ca^2+^ efflux by ∼ 3-fold. In contrast, extracellular Na^+^ presence or absence in KO-derived hepatocytes failed to enhance Ca^2+^ efflux pathway in either system. Moreover, Na^+^ influx via the cell membrane is required for full activation of the mitochondrial Na^+^/Ca^2+^ exchanger, this is consistent with reports demonstrating a low affinity of NCLX for cytosolic Na^+^ (4, 29).

What is the basis for the differences in the present and previous studies that supported a Na^+^-independent primary mechanism for mitochondrial Ca^2+^ efflux in hepatocytes (14)? Many of these studies were conducted on isolated mitochondria that often lose their mitochondrial membrane potential following their isolation. However, even a small drop of mitochondrial ΔΨ is sufficient to inhibit Na^+^-dependent mitochondrial Ca^2+^ efflux by NCLX (24, 30). Interestingly, isolated liver mitochondria from energized hepatocytes, treated with a beta-adrenergic receptor agonist, restored the Na^+^-dependent Ca^2+^ efflux system (22). Thus, if the isolated mitochondria were not fully energized, mitochondrial Na^+^-dependent Ca^2+^ efflux could have been impaired. A Na^+^-independent mitochondrial Ca^2+^ efflux was also suggested in a study conducted on cultured hepatocytes, which compared their mitochondrial Ca^2+^ efflux to cardiomyocytes and neuronal cells (15). The reason for the differences between this and our study are unclear, however, the former study was done on primary hepatocytes 7 days post their culturing, a timeline which may lead to hepatocytes de-differentiation (31). Moreover, mitochondrial Ca^2+^ efflux in this study was exceedingly 100-fold slower than rates found in previous reports and our studies, indicating that the mitochondrial Ca^2+^ efflux was largely inactive (15).

Hepatic stimulation by glucagon and other hepatic hormones, at low physiological concentrations that are sufficient to facilitate Ca^2+^ release from the ER stores into the cytoplasm, evoke low frequency cytosolic Ca^2+^ oscillation spikes. Importantly, these spikes are transmitted individually into the mitochondrial matrix enhancing the metabolic activity (9, 10, 26). Our results show that while deletion of hepatic NCLX has a subtle effect on glucagon and vasopressin induced cytosolic oscillations, it triggers a profound inhibition of glucagon-dependent mitochondrial Ca^2+^ oscillations. What is the molecular basis for this effect of NCLX on mitochondrial Ca^2+^ oscillations? Previous studies indicated that mitochondrial Ca^2+^ influx by MCU is downregulated by even a subtle rise in mitochondrial free Ca^2+^, a process that is tuned by EMRE and Micu1 (32–35). Consistent with these findings, mitochondrial Ca^2+^ oscillations are triggered by caffeine in neurons and are suppressed by NCLX KO, which causes mitochondrial Ca^2+^ overload (36).

The role of Ca^2+^ signaling in controlling metabolic activity and gluconeogenesis is of intense interest (13). Studies using the hepatic hormones or other agonists often evoked a strong single cytosolic and mitochondrial Ca^2+^ response and not the hallmark oscillatory response encountered during physiological activity (12, 13). Our results show that impairing the Ca^2+^ oscillatory pattern activated by glucagon and VP in the KO model of NCLX is sufficient to affect both the metabolic activity and gluconeogenesis. Another issue that is not resolved, is the role of the hormone-dependent cytosolic or mitochondrial Ca^2+^ response in hepatic gluconeogenesis (26).

While a mitochondrial Ca^2+^ rise is linked to metabolic regulation, many other studies underscore the importance of extra mitochondrial pathways in mediating the Ca^2+^ dependent metabolic changes, most notably the Aralar pathway in neurons and cardiomyocytes (37). The hepatic KO of NCLX offers a unique paradigm to address this issue because in using this model the hormone dependent cytosolic Ca^2+^ oscillations are fully preserved while the mitochondria are suppressed. Our results indicate that the suppression of the mitochondrial Ca^2+^ oscillations are sufficient to reduce the glucagon dependent mitochondrial metabolic activity and gluconeogenesis thus underscoring the dominant role of mitochondrial Ca^2+^ signaling in this process.

Collectively, the results of this study reveal that NCLX plays an essential role in regulating hepatic mitochondrial calcium extrusion in a Na^+^-dependent route. It underscores the physiological evidence of NCLX in regulating mitochondrial Ca^2+^ oscillations required for efficient electron transport chain and oxidative phosphorylation. Lastly, this study, sheds light on a metabolic downstream glucagon-mediated Ca^2+^ signaling role of NCLX, which by facilitating the transmission of cytosolic Ca^2+^ oscillations to mitochondria drives the transcription-independent, gluconeogenesis pathway. Tuning NCLX activity may open future avenues for targeted therapeutic strategies to overcome hepatic metabolic disorders, such as nonalcoholic fatty liver disease or type-2 diabetes.

## Methods

### Animal models

Wildtype C57BL/6NJ mice and NCLX-null (C57BL/6NJ-*Slc8b1^em1(IMPC)J^*/J) were purchased from Jackson laboratories (Jackson lab, Bar Harbor, ME), and bred in our vivarium. Due to global loss of NCLX from birth in these mice, their use was restricted for in vitro experiments and for validation purposes in a different NCLX-null model. Mice genotyping was performed on earpieces or clipped tails obtained during the weaning of pups. Genotyping was performed following the protocol of Jax laboratories by real-time polymerase chain reaction. The following primers were used to detect NCLX-null^−/−^, Heterozygous^+/−^, or WT^+/+^ mice: Forward primer-GGCTCCTGTCTTCCTCTGTG and Reverse primer-GTGTCCATGGGCTTTTGTG.

In addition, a conditional liver-specific NCLX knockout, NCLX floxed mice containing loxP sites flanking exons 5-7 of the *Slc8b1* gene (ch12: 113298759-113359493) was generously provided by Prof. John Elrod (21). Floxed mice were injected via the tail vein with an AAV8-Cre under the control of a hepatocyte-specific promoter TBG; (1.3 × 10^11^ plaque-forming units per mouse, ((107787-AAV8, Addgene, Watertown, MA)). AAV8-TBG-Null (105536-AAV8, Addgene) was injected in floxed littermates of the control group. Animals were used for experiments 5 weeks post injection.

Experimental procedures conducted on mice were performed in accordance with animal welfare and in compliance with other related ethical regulations. The mice studies were conducted under an approved Institutional Animal Care and Use Committee (IACUC) protocol the Ben-Gurion University. The mice were fed standard chow diet and maintained under controlled conditions (housing at 22 °C with a 12:12 h light:dark cycle). For selected experiments, mice were fasted with free access to water and were compared with ad libitum mice. For in vivo experiments and analysis, a randomization and a double blinded-manner were performed using ear-tagging and random mice numbering systems which were revealed after the termination of the experiment.

### Hepatocytes isolation

Hepatocytes were isolated from adult male mice by a two-step collagenase perfusion method as previously reported (26, 35, 38). In brief, the liver was perfused through abdominal inferior vena cava and hepatic portal vein was cut through. The organ was washed with Hanks’ calcium and magnesium-free buffer for 6 min After the liver had been freed of blood, the calcium-free buffer was replaced by the liver digest media (17703-034, Thermo Fisher Scientific, Roskilde, Denmark) for 7 min. A perfusion rate of 5 ml/min and a temperature of 37°C were maintained for both solutions during the entire procedure. After the perfusion, the gallbladder and remnants of the diaphragm were removed, and the liver was rapidly excised from the body cavity and transferred to a sterile Petri dish containing plating medium. The cells were a filtered by 70-micron strainer. Hepatocytes were then gently washed by low-speed centrifugation at 50 g for 5 min at 4°C. Cells were then diluted with Percoll (P1644, Sigma-Aldrich) at ratio 1:1 and centrifuged for 10 min and the pellet was collected. Hepatocyte viability and yield were determined by trypan blue exclusion, with isolations only over 70% viability being used. Primary hepatocytes were resuspended and plated in “plating media”: (DMEM (11965092, Thermo) supplemented with 10% Fetal Bovine serum (F2442, Sigma-Aldrich), 100 U/ml penicillin and 100 μg/ml streptomycin (03-031-1B, Biological industries), 2mM Sodium pyruvate (113-24-6, Sigma-Aldrich),1 µM dexamethasone (SC-29059, Santa Cruz) and100 nM insulin (Actrapid, Novo Nordisk)). For imaging experiments, hepatocytes were plated directly on collagen precoated coverslips at a density of 1 million cells per 20 cm2; for seahorse respirometry assays 8000 hepatocytes were plated per well and for biochemical assays cells were plated at a density of one cell per well in six-wells plate. Two hours after hepatocytes seeding, media was replaced to remove unattached hepatocytes with a “maintenance medium” containing: DMEM medium supplemented with 100 U/ml penicillin and 100 μg/ml streptomycin, 2mM Sodium pyruvate, 100 nM dexamethasone, 1nM Insulin and 0.2% Fraction V fatty-acid free bovine serum albumin (BSA, E588, VWR, Radnor, PA). Cells were kept for 24-36 h.

### Cell lines

Clonal HepG2 cells were cultured in low glucose (5 mM) Dulbecco’s Modified Eagle Medium (MFCD00217342, Sigma-Aldrich, St. Louis, MO) supplemented with 10% FBS, 50 U/ml penicillin, and 5 mM HEPES buffer. For imaging experiments, HepG2 cells were plated seeded onto collagen coated coverslips and Ca^2+^ imaging was performed 72 h post seeding.

### Fluorometry for liver isolated mitochondria

The livers were perfused with Krebs-Henseleit bicarbonate buffer containing (in mM): 120 NaCl, 4.8 KCl, 1.2 MgSO_4_, 1.2 KH_2_PO_4_, 1.3 CaCl2, 25.3 NaHCO3 and or 10mM-Tris/lactate plus 1 mM-Tris/pyruvate, pH 7.4) supplemented with norepinephrine (1 µM) before isolating the mitochondria as described in Crompton studies (Goldstone, et al. 1983).

Mitochondria from liver were then isolated as previously described (Renault, et al. 2016). Briefly C57BL/6N mice (1-2 months) were anesthetized with isoflurane, liver was removed immediately and sliced into Trehalose isolation buffer (TIB) buffer containing: 270mM sucrose, 10mM HEPES-KOH (pH=7.4), 10mM KCl, 1mM EDTA, 0.1% BSA and a freshly protease inhibitor was supplemented while kept on ice all the time. The liver was mechanically homogenized using 10 strokes using a Teflon glass homogenizer. Then a set of centrifugations was performed to obtain the mitochondria fractions. All the steps were performed in a swinging bucket centrifuge at 4C. The homogenate was first centrifuged at 600 g for 10 min to remove nuclei and cell debris. The supernatant was transferred to a new conic tube and centrifuged at 3500 g for 10 min. The resultant pellet was resuspended in TIB buffer and re-centrifuged at 1500 g for 5 min. The supernatant was transferred again to a new conic tube and centrifuged at 5500 g for 10 min. The pellet was then collected and resuspended with TIB buffer and saved on ice as the mitochondrial fraction. Protein concentration was determined by Bradford assay (500-0006, Bio-Rad, Hercules, California, US).

To measure mitochondrial calcium uptake and efflux, Extra-mitochondrial Ca^2+^dye Oregon-Green (Thermo) (0.5 µM) was added to mitochondria resuspended in a Ca^2+^-free TIB buffer in a cuvette at a protein concentration of 1mg/1ml. Fluorescence was measured with constant stirring, in a cuvette fluorimeter at 496 nm excitation and 524 nm emission, as previously described (25). Measurements started from baseline followed by the addition of Ca^2+^ bolus (6 µM) to trigger the mitochondrial calcium influx followed by addition of ruthenium red (10 µM) in the presence of sodium (18 mM) or NMDG (18 mM) to monitor the mitochondrial calcium efflux phase rates.

### Immunoblotting

Isolated organ tissues were washed in ice-cold PBS, minced, and homogenized in MAS buffer (70 mM sucrose, 220 mM mannitol, 5 mM KH_2_PO_4_, 5 mM MgCl_2_, 1 mM EGTA, 2 mM HEPES pH 7.4). Liver, Heart and lung were mechanically homogenized with 10-15 strokes in glass-glass Teflon-glass homogenizer until all tissues were solubilized. All homogenates were centrifuged at 1,000 *g* for 10 min at 4°C; then, the supernatant was collected. Protein concentration was determined by Bradford assay (Bio-Rad). Twenty micrograms protein of tissue homogenates were mixed with 4× LDS sample buffer and subjected to SDS-PAGE gel electrophoresis. Using a wet transfer system (Bio-Rad). The membranes were blocked with 5% nonfat dry milk for 1 h and then incubated with the following antibodies: NCLX (1:1000, sc-161921, Santa Cruz), MCU (1:1000, sc-515930, Santa Cruz) and VDAC1 (1:500, sc-390996, Santa Cruz). Antibodies were used according to the manufacturer’s instructions. After overnight incubation, membranes were washed with Tris-buffered saline containing 0.1% Tween-20 and then incubated with anti-rabbit IgG secondary antibody coupled to HRP (1:10000, sc-2357, Santa Cruz) for 1 h or an anti-mouse (1:5000, sc-2005, Santa Cruz) solution for 1 h. The membranes were washed again as mentioned above and then exposed to a chemiluminescent protein detection system (Fusion SOLO X, Vilber). Detection was done with the EZ-ECL Chemiluminescence Detection kit for HRP (20-500-120, Sartorius).

### Live fluorescence imaging

Kinetic live-cell fluorescent imaging was performed to monitor Ca^2+^ transients using two imaging systems. The first system consisted of an Axiovert 100 inverted microscope (Zeiss, Oberaue, Germany), Polychrome V monochromator (Till Photonics, Planegg, Germany), and a Sensi-Cam cooled charge-coupled device (PCO, Kelheim, Germany). Fluorescence images were acquired with Imaging WorkBench 6.0 software (Axon Instruments, Foster City, CA, USA). The second system consisted of an IX73 inverted microscope (Olympus) equipped with pE-4000 LED light source and Retiga 600 CCD camera. All images were acquired through a ×20/0.5 Zeiss Epiplan Neofluar objective using Olympus cellSens Dimension software.

Cells were pre-loaded with the indicated calcium specific fluorescent dye at the indicated concentrations for 30 min at 37°C using a modified Krebs-Ringer’s solution containing (in mM): 126 NaCl, 5.4 KCl, 0.8 MgCl2, 20 HEPES, 1.8 CaCl2, 15 glucose, 10 lactate, 1 pyruvate with pH adjusted to 7.4 with NaOH and supplemented with 0.1% BSA. After dye loading, cells were washed three times with fresh dye-free Krebs-Ringer’s solution, followed by additional incubation of 30 min to allow for the de-esterification of the residual dye.

For mitochondrial Ca^2+^ measurements [Ca^2+^]m, cells were loaded with 1 μM Ca^2+^-specific dye Rhod2-AM (50024, Biotium, Fremont, CA) that preferentially localizes in mitochondria. Rhod2-AM was excited at 552 nm wavelength light and imaged with a 570 nm-long pass filter as previously described (Kostic, et al. 2015).

For cytosolic Ca^2+^ measurements [Ca^2+^]c, cells were loaded either with Fura-2AM ratiometric dye (2 μM) (0102, TEF Labs and F0888, Sigma-Aldrich) or Flou4-AM (2μM) (1892, Lumiprobe, Cockeysville, MD). Fura-2AM was excited alternately with 340 nm and 380 nm excitations and imaged with a 510 nm long pass filter, as described previously ((29, 39)). Flou4-AM was excited at 494 nm and imaged at 506 nm.

At the beginning of each experiment, cells were perfused with Ca^2+^-containing Krebs-Ringer’s solution supplemented with 0.1% fatty-acid free BSA (E588, VWR). To trigger ionic responses, cells were perfused with supplemented Krebs-Ringer’s containing ATP (5μM) (0220, Amresco), Vasopressin (1nM) (sc-356188, Santa Cruz), Glucagon (100nM). (16941-32-5, Sigma-Aldrich).

Traces of Ca^2+^ responses were analyzed and plotted using KaleidaGraph. Oscillatory waves frequencies and area under the curve (AUC) of each graph were calculated using GraphPad Prism 10. The rate of ion transport was calculated from each graph (summarizing an individual experiment) by a linear fit of the change in the fluorescence (Δ*F*) for Ca^2+^ influx and efflux over time (Δ*F*/d*t*). Rates from *n* experiments (as mentioned in legends to the figures) were averaged and displayed in bar graph.

Mitochondrial mass was assessed by staining primary hepatocytes with 200 nM MitoTracker™ Deep Red FM (MTDR) (M22426, Thermo) for 1 h. Cells were washed with PBS 3 times and Tecan Spark 10M multimode plate reader was utilized to read fluorescence intensity. MTDR was excited with 633 nm wavelength and emission collected at 665 nm.

### Digitonin-permeabilized hepatocytes

To determine the Na+ dependence of mitochondrial Ca^2+^ efflux, a digitonin-permeabilized cell system was assayed as previously described with modifications detailed below (4). In brief, isolated hepatocytes were plated on collagen coated coverslips, the hepatocytes, were washed three times with PBS and loaded with a Rhod2-AM dye (1 µM) and maintained in a sucrose buffer containing (in mM): 220 Sucrose, 10 HEPES, 5 Succinate, 2.5 KH_2_PO_4_, 0.4 EGTA with the pH adjusted to 7.4 with KOH. During incubation time Cyclosporine A (1 µM) was supplemented to the media. Coverslips were first perfused with 4µg/ml digitonin-supplemented sucrose buffer (D180-1, Goldbio) for 30 s, then the coverslips were then washed with digitonin-free buffer. Measurements were started with perfusion of Ca^2+^-free buffer to maintain a baseline, followed by the addition of Ca^2+^ bolus (6 µM). After reaching maximal Ca^2+^uptake, 10 µM of RR was added to block Ca^2+^ uptake in a solution containing either 10mM of Na^+^ or alternatively NMDG^+^.

### Biochemical assessments

For Gluconeogenesis and hepatic glucose production, assays were performed as previously described (40). In brief, primary mouse hepatocytes were isolated and seeded in six-well plates in a 1 × 10^6^ cell per well density and kept in maintenance media. 24 h later, Cells were washed with PBS 3 times and incubated with 1 ml of glucose-free DMEM without phenol red, supplemented with 20 mM sodium lactate and 2 mM sodium pyruvate, 4mM glutamine, 100nM dexamethasone and supplemented with fatty-acid free BSA 0.1%. The media contained one of the three treatments: Glucagon (20nM), Glucagon and Insulin (20nM and 100 nM respectively) or vehicle. After an 8 h incubation, 200 μl of the medium was collected and pelleted at slow centrifugation to remove floating cells or cell debris. 30 μl of the supernatants was taken to measure glucose using a colorimetric glucose oxidase assay kit (A22189, Thermo). Glucose values were normalized to the amount of proteins for each individual well. For Pyruvate carboxylase activity assessment assays, an ELISA based assay was used (K2075, Abcam, Cambridge, MA) according to the manufacturer instructions.

### Respirometry assays

Respirometry of primary mouse hepatocytes were performed using the Seahorse Bioscience XFe96 platform (Agilent Technologies, Santa Clara, CA) as previously described (40). In brief, primary isolated hepatocytes were plated at 8,000 cells/well in a collagen coated XFe96 plate. Cells were cultured overnight in plating medium used in the isolation. The next day, growth media was replaced to glucose-free media supplemented with 10:1 mM lactate to pyruvate and cells were treated with glucagon (20 nM) and or insulin (100 nM) for 8 h incubation. Medium was then replaced with XF DMEM media, pH 7.4 supplemented with glutamine, lactate and sodium pyruvate. Mitochondrial stress tests were performed via sequential injection of 3 μM oligomycin, 3 μM trifluoromethoxy carbonyl-cyanide phenylhydrazone (FCCP), and 2μM rotenone/antimycin A. OCR was normalized by cell count in each well. Cells were fixed with 4% paraformaldehyde (J61899, Thermo) and normalized by cell count. Cells were stained with 1 μg/ml DAPI (D1306, Thermo). A Tecan Spark 10M multimode plate reader was utilized to read fluorescence intensity at ex/em (358/461).

### Statistical analysis

GraphPad Prism 10 software was used for statistical analysis and Adobe Illustrator for the graphic design of the figures. Statistical significance was assessed by two-tailed unpaired Student’s *t*-test for two-group comparison and one-way or two-way analyses of variance (ANOVAs) grouped analyses followed by Tukey’s test for multiple-group comparisons. All bar graphs were presented as averaged individual sample values of *n* measurements ± SEM. A value of p < 0.05 accepted as statistically significant. Symbols of significance used: ns (non-significant), p > 0.05, * p ≤ 0.05, ** p ≤ 0.01, *** p ≤ 0.001.

## Acknowledgements

The authors would like to thank those who contributed helpful discussions, insight and support of the research, including Dr. Marc Liesa-Roig and Dr. Eli Lewis and members of the Sekler lab. This work was supported by Israeli science foundation ISF (1424/17, DIP SE 2372/1-1) grants. E.A.A. was an Azrieli foundation fellow.

## Author Contributions

Conceptualization, E.A.A., M.T. and I.S..; Methodology, M.T., E.A.A., M.K. and I.S.; Investigation, M.T. and E.A.A.; Writing –Original Draft, E.A.A, M.T. and I.S.; Writing–Review & Editing, E.A.A., M.T., M.H., G.E.S., O.S.S and I.S..; Funding Acquisition, I.S.; Resources, M.H. and I.S. Supervision, E.A.A., M.T., O.S.S, M.H. and I.S. All authors read and approved the final version of the manuscript.

## Supporting Information

**Fig. S1.**
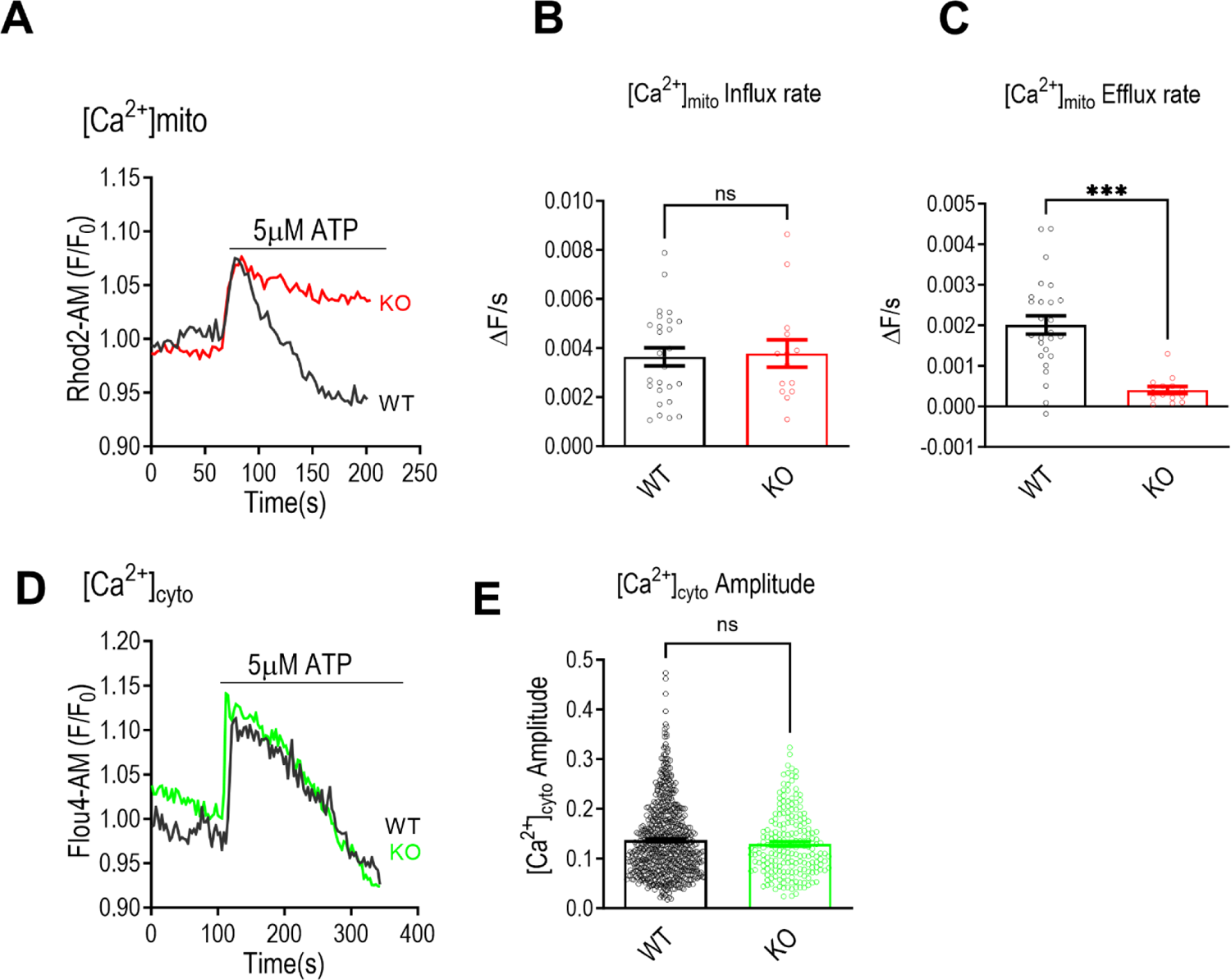
(A-C) Mitochondrial Ca^2+^ kinetics evoked by ATP in cultured primary hepatocytes isolated from global NCLX KO mice and their WT littermates. (A) Representative mitochondrial Ca^2+^ transients of NCLX KO hepatocytes and their WT controls, triggered by ATP (5μM). Mitochondrial Ca^2+^ was monitored by the mitochondrial Ca^2+^ dye, Rhod2-AM. (B,C) Quantification of [Ca^2+^]m influx and efflux rates (as in A) for WT (n = 26) and NCLX KO (n = 14). (D,E) Cytosolic Ca^2+^ kinetics evoked by ATP in cultured primary hepatocytes isolated from NCLX KO mice and their WT littermates. (D) Representative cytosolic Ca^2+^ transients of NCLX KO hepatocytes and their WT controls, triggered by ATP (5μM). Cytosolic Ca^2+^ was monitored by Flou4-AM. (E) Quantification of [Ca^2+^]c amplitude (A) for WT (n = 640) and NCLX KO (n = 204). All values are represented as mean ± SEM, \**p* < 0.05; \*\**p* < 0.01; \*\*\*\**p* < 0.0001, N.S-non-significant (two-tailed unpaired *t*-test for comparisons between two groups was used and one-way ANOVA with Tukey’s multiple comparison test was used for three or more groups).

**Fig. S2.**
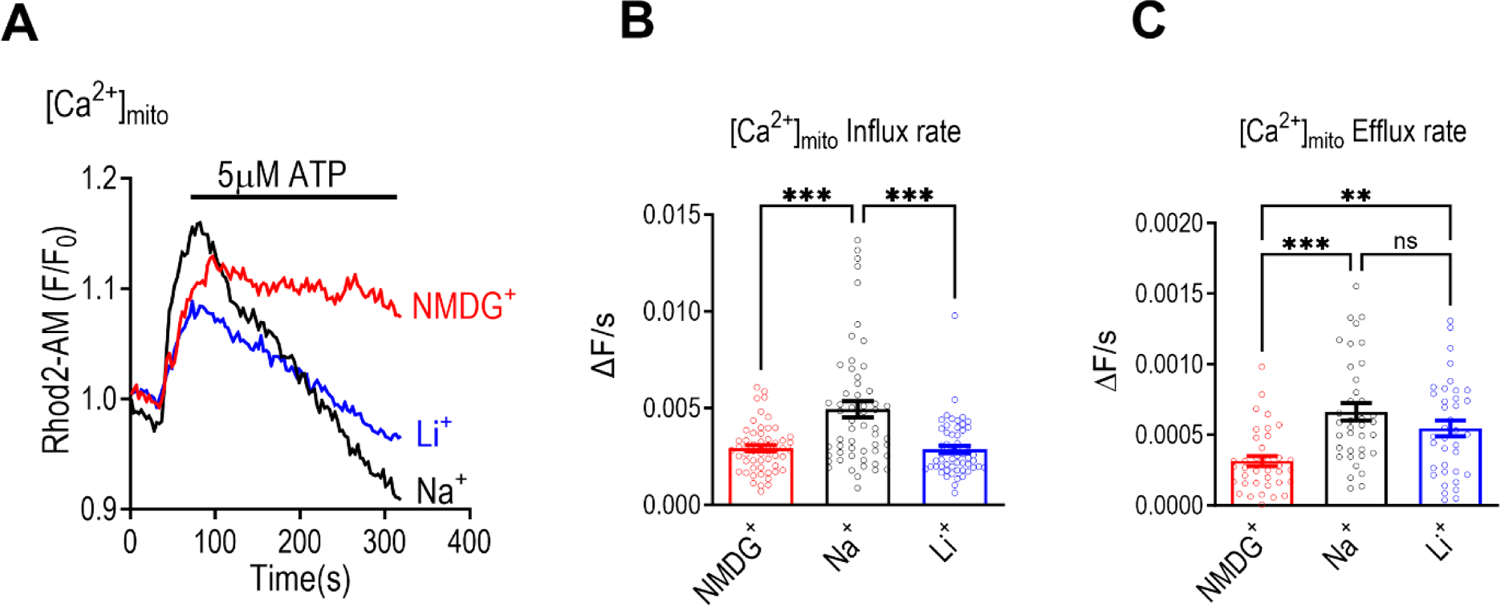
(A-C) Analysis of monovalent cation dependence of mitochondrial Ca^2+^ efflux in human HepG2 cells. (A) Representative traces of mitochondrial Ca^2+^ transients upon application of ATP to human HepG2 cells in the presence of Na^+^, Li^+^ or NMDG^+^ (iso-osmotically replacing Na^+)^. (B) Quantification of [Ca^2+^]m influx rates in HepG2 cells upon ATP stimulation (n = 56 for all conditions). (C) Quantification of [Ca^2+^]m efflux rates in HepG2 cells upon ATP stimulation (n = 37 for all conditions). All values are represented as mean ± SEM, \**p* < 0.05; \*\**p* < 0.01; \*\*\*\**p* < 0.0001, N.S-non-significant (two-tailed unpaired *t*-test for comparisons between two groups was used and one-way ANOVA with Tukey’s multiple comparison test was used for three or more groups).

**Fig. S3.**
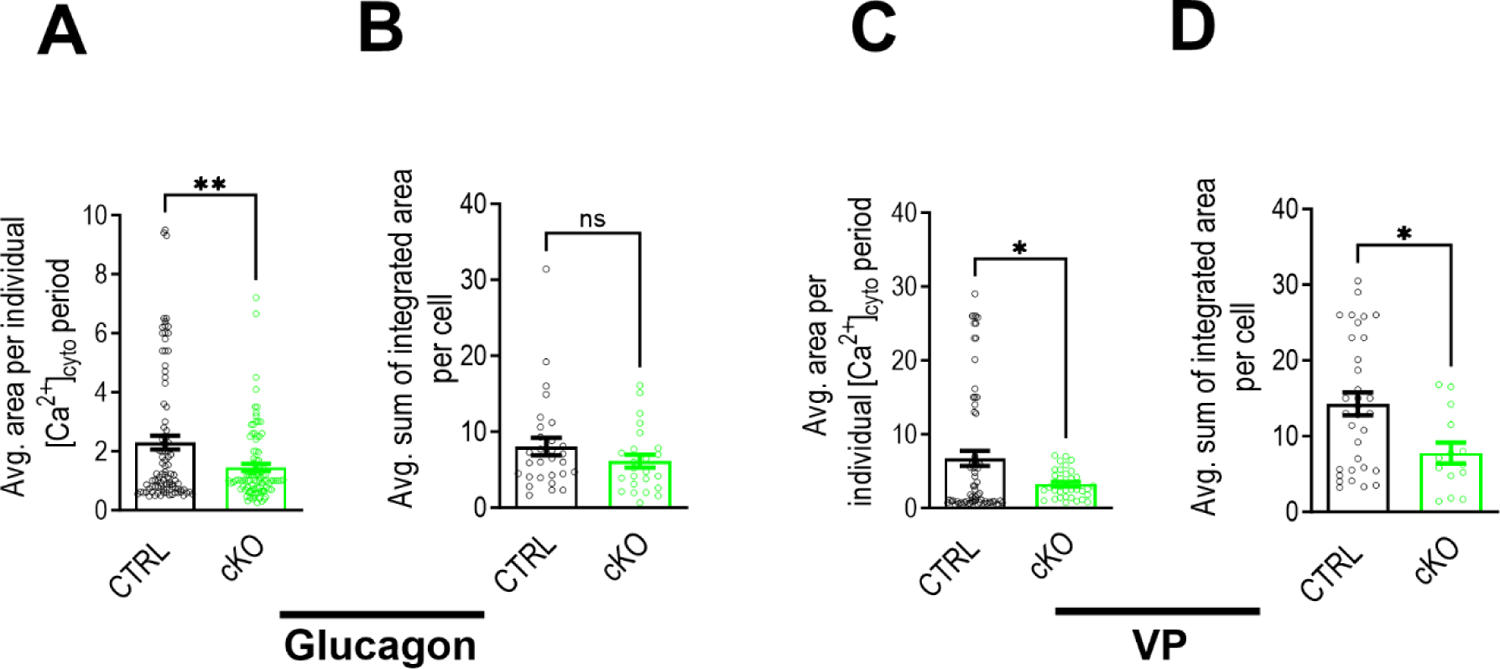
(A.B) Analysis of cytosolic Ca^2+^ oscillations (see figure 3) triggered by physiological concentrations of glucagon in cultured primary hepatocytes isolated from NCLX cKO mice and their controls. (A) Quantification of average area under curve per individual period of [Ca^2+^]c wave shown in for CTRL (n = 94) and NCLX cKO hepatocytes (n = 101). (B) Quantification of average sum of integrated area under curve of [Ca^2+^]c waves per cell shown in for CTRL (n = 29) and NCLX cKO hepatocytes (n = 24). (C.D) Analysis of Cytosolic Ca^2+^ oscillations triggered by physiological concentrations of VP in cultured primary hepatocytes isolated from NCLX cKO mice and their controls. (C) Quantification of average area under curve per individual period of [Ca^2+^]c wave shown in for CTRL (n = 70) and NCLX cKO hepatocytes (n = 35). (D) Quantification of average sum of integrated area under curve of [Ca^2+^]c waves per cell shown in for CTRL (n = 33) and cKO hepatocytes (n = 14). All values are represented as mean ± SEM, \**p* < 0.05; \*\**p* < 0.01, N.S-non-significant (two-tailed unpaired *t*-test for comparisons between two groups was used and one-way ANOVA with Tukey’s multiple comparison test was used for three or more groups).

**Fig. S4.**
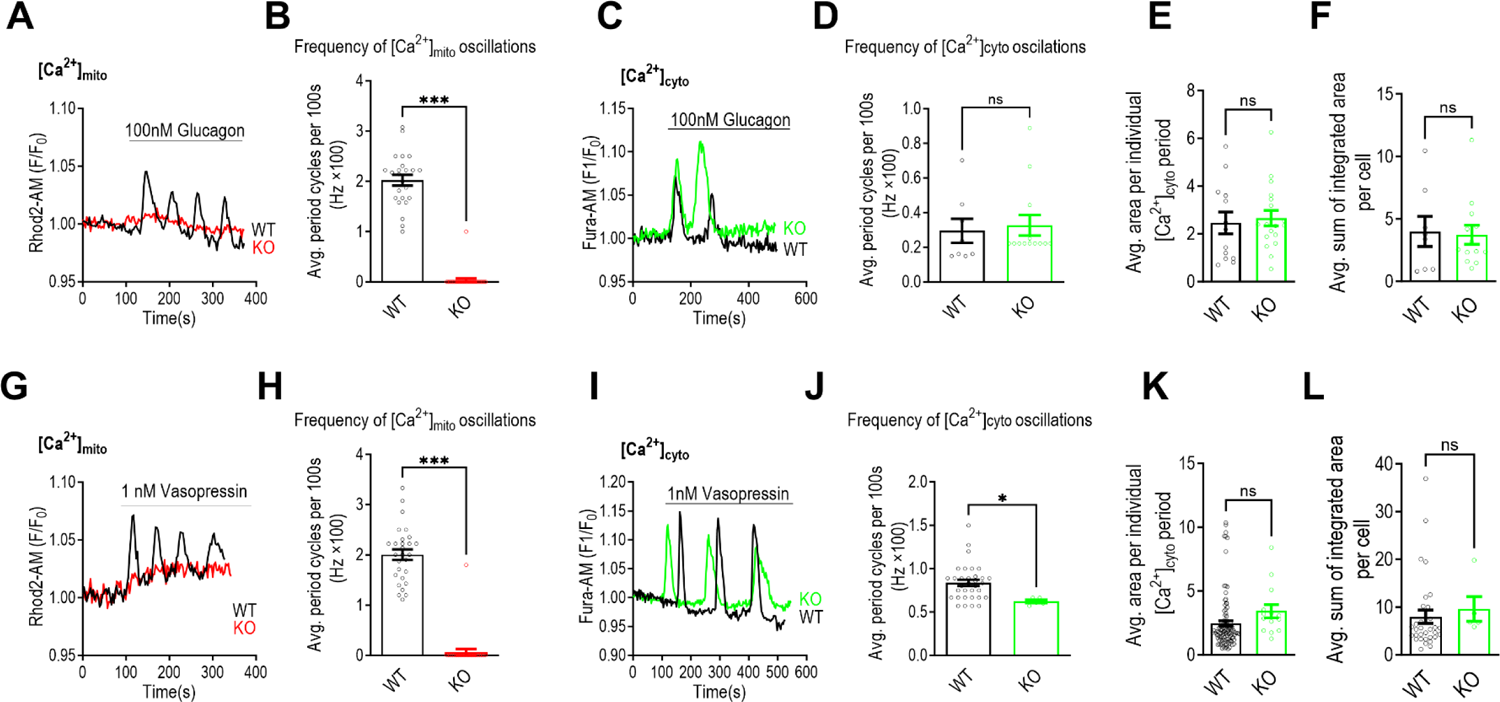
(A,B) mitochondrial Ca^2+^ oscillations triggered by physiological concentrations of glucagon in cultured primary hepatocytes isolated from global NCLX KO (Red) and their WT controls (black). Mitochondrial Ca^2+^ was monitored by Rhod2-AM. (A) Representative fluorescence traces of mitochondrial Ca^2+^ oscillations triggered by glucagon (100nM) in NCLX KO hepatocytes and their WT controls. (B) Quantification of [Ca^2+^]m frequency of oscillations shown in (A) for WT (n = 24) and NCLX KO hepatocytes (n = 29). (C-F) Cytosolic Ca^2+^ oscillations triggered by physiological concentrations of glucagon in cultured primary hepatocytes isolated from NCLX KO mice (Green) and their WT controls (black). Cytosolic Ca^2+^ was monitored by Fura2-AM. (C) Representative fluorescence traces of cytosolic Ca^2+^ oscillations triggered by glucagon (100 nM) in NCLX KO hepatocytes and their WT controls. (D) Quantification of [Ca^2+^]c frequency of oscillations shown in (C) for WT (n = 8) and NCLX KO hepatocytes (n = 13). (E) Quantification of average area under curve per individual period of [Ca^2+^]c wave shown in for WT (n = 13) and NCLX KO hepatocytes (n = 18). (F) Quantification of average sum of integrated area under curve of [Ca^2+^]c waves per cell shown in for WT (n = 8) and NCLX KO hepatocytes (n = 13). (G,H) Mitochondrial Ca^2+^ oscillations triggered by physiological concentrations of vasopressin (VP) in cultured primary hepatocytes isolated from NCLX KO mice (Red) and their WT controls (black). Mitochondrial Ca^2+^ was monitored by Rhod2-AM. (G) Representative fluorescence traces of mitochondrial Ca^2+^ oscillations triggered by VP (1nM) in NCLX KO hepatocytes and their WT controls. (H) Quantification of [Ca^2+^]m frequency of oscillations shown in (G) for WT (n = 28) and NCLX KO hepatocytes (n = 28). (I-L) Cytosolic Ca^2+^ oscillations triggered by physiological concentrations of VP in cultured primary hepatocytes isolated from NCLX KO mice (Green) and their WT controls (black). Cytosolic Ca^2+^ was monitored by Fura2-AM. (I) Representative fluorescence traces of cytosolic Ca^2+^ oscillations triggered by VP (1 nM) in NCLX KO hepatocytes and their WT controls. (J) Quantification of [Ca^2+^]c frequency of oscillations shown in (A) for WT (n = 33) and NCLX KO hepatocytes (n = 5). (K) Quantification of average area under curve per individual period of [Ca^2+^]c wave shown in for WT (n = 104) and NCLX KO hepatocytes (n = 14). (L) Quantification of average sum of integrated area under curve of [Ca^2+^]c waves per cell shown in for WT (n = 32) and NCLX KO hepatocytes (n = 5). All values are represented as mean ± SEM, \**p* < 0.05; \*\**p* < 0.01; \*\*\*\**p* < 0.0001, N.S-non-significant (two-tailed unpaired *t*-test for comparisons between two groups was used).

